# Key role of piRNAs in telomeric chromatin maintenance and telomere nuclear positioning in *Drosophila* germline

**DOI:** 10.1101/163030

**Authors:** Elizaveta Radion, Valeriya Morgunova, Sergei Ryazansky, Natalia Akulenko, Sergey Lavrov, Yuri Abramov, Pavel A. Komarov, Sergey I. Glukhov, Ivan Olovnikov, Alla Kalmykova

## Abstract

Telomeric small RNAs related to PIWI-interacting RNAs (piRNAs) were discovered in different species, however, their role in germline-specific telomere function remains poorly understood. Using a *Drosophila* model, we show that the piRNA pathway provides a strong germline-specific mechanism of telomere homeostasis. We show that telomeric retrotransposon arrays belong to a unique class of dual-strand piRNA clusters whose transcripts, required for telomere elongation, serve simultaneously as piRNA precursors and their only targets. However, the ability to produce piRNAs and bind Rhino – a germline-specific homolog of heterochromatic protein 1 (HP1) – varies along telomeres. Most likely, this heterogeneity is determined by the peculiarities of telomeric retrotransposons themselves. piRNAs play a pivotal role in the establishment and maintenance of telomeric and subtelomeric chromatin in the germline facilitating loading of HP1 and histone 3 lysine 9 trimethylation mark – highly conservative telomere components – at different telomeric regions. piRNA pathway disruption results in telomere dysfunction characterized by a loss of heterochromatic components and translocation of telomeres from the periphery to the nuclear interior but does not affect the telomere end capping.

## INTRODUCTION

Telomere transcription is an evolutionary conserved feature of eukaryotic telomeres (1). Biogenesis of telomeric transcripts has been shown to be tightly connected to telomere length control and telomeric chromatin formation. Telomeric transcripts serve as precursors for small RNAs (tel-sRNAs) discovered in mammalian embryonic stem cells, in the ciliate *Tetrahymena thermophila*, in plants, and in *Diptera* (2-5). Small RNAs generated by the subtelomeric regions in fission yeast as well as some tel-sRNAs have been implicated in the assembly of telomeric heterochromatin (2, 5, 6). Plant and mammalian tel-sRNAs are related to the class of Piwi-interacting RNAs (piRNAs) generated in germline and in stem cells (7, 8). However, the role of tel-sRNAs in the germline-specific telomere function is poorly understood. Telomeric piRNAs and their role in telomere length control was first described in *Drosophila melanogaster* (3). Using a *Drosophila* model, we performed in-depth study of the biogenesis and function of telomeric piRNAs in the germline.

The piRNA-mediated pathway provides silencing of transposable elements (TE) in the germline (7,9). In contrast to small interfering RNAs (siRNA), which are processed by the Dicer endonuclease from double strand RNA, the piRNAs are generated from long single strand precursor transcripts. These piRNA precursors are encoded by distinct genomic regions enriched by damaged TE copies termed piRNA clusters (10). The dual-strand piRNA clusters found in the *Drosophila* germline produce piRNAs from precursors transcribed by both genomic strands. The dual-strand piRNA clusters can be classified into several types such as extended pericentromeric TE-enriched regions (10), individual TE-associated euchromatic clusters (11), and transgene-associated clusters (12, 13). In all these instances, piRNAs and their targets are expressed by distinct genomic regions.

These pericentromeric piRNA clusters produce piRNAs that target the nascent transcripts of the active TEs resulting in the posttranscriptional silencing as well as heterochromatin protein 1 (HP1)-mediated transcriptional silencing (14-17). In the perinuclear compartment, the piRNA-mediated cleavage of TE transcripts initiates further processing of the cleavage products to increase piRNA abundance and diversity (10, 18-20). Distinct chromatin components of the piRNA clusters that couple transcription and RNA transport appear to direct the cluster-derived transcripts into the piRNA processing machinery (21-23). The germline-specific homolog of HP1 – Rhino (Rhi) – is essential for piRNA production from the dual-strand piRNA clusters (24-26). piRNAs are required at early embryonic stages for deposition of the Rhi and Histone 3 lysine 9 trimethylation mark (H3K9me3) at dual-strand piRNA clusters, but at later developmental stages the chromatin of piRNA clusters is maintained by an unknown Piwi-independent mechanism (27).

The *Drosophila* telomeres produce abundant piRNA and appear to constitute a separate class of piRNA clusters. The telomeres of *D. melanogaster* are maintained by transpositions of the specialized telomeric retrotransposons, while the telomerase gene has likely been lost in an ancestor of *Diptera* (28). The non-LTR *HeT-A, TART,* and *TAHRE* retroelements are organized in tandem head-to-tail telomeric arrays with *HeT-A* being the prevailing telomeric retrotransposon (29-31). The telomere associated sequences (TAS) consist of complex satellite-like repeats and are located proximally to retrotransposon arrays. Telomeric transcripts are processed into piRNAs that regulate telomeric TE expression, and their transposition rate onto chromosome ends in the germline (3, 32).

Thus, the telomeres need to produce both the intact TE mRNA required for telomere elongation, as well as piRNAs that target these TE mRNA in order to regulate optimal telomere length. Indeed, altering this balance can lead to disruption of telomere length control (3, 33). Different factors including the piRNA pathway components, act cooperatively to regulate telomeric repeat expression and telomere protection in the germline providing genome integrity during early development (34, 35). It is clear, that the involvement of telomeres in piRNA production should considerably affect telomere biology in the germline, however, telomeres have not yet been characterized as piRNA clusters.

Analysis of ovarian small RNA-seq data revealed abundant piRNAs corresponding to both genomic strands of telomeric retrotransposons and TAS (10), thus telomeric piRNA clusters can be formally related to the dual-strand type. However, the main distinction of the telomeric piRNA clusters is that their transcripts serve both as a source of piRNAs and as their only targets. While genome-wide data argues against the existence of defined promoters at dual-strand clusters (36, 37), the telomeric retroelements are characterized by the presence of bidirectional promoters that provide transcription of the piRNA precursors (38-40).

Experimental evidence indicates that *HeT-A* and related *TAHRE* elements are extremely sensitive to piRNA pathway disruption showing up to 1,000-fold overexpression in contrast to *TART*, which demonstrates only modest upregulation (3, 32, 41, 42). Therefore, *HeT-A* expression has been extensively used as a readout of piRNA pathway disruption. However, *HeT-A* elements are not typical piRNA targets, but they belong to piRNA clusters. In contrast to *HeT-A*, the expression of germline piRNA clusters even decreased upon disruption of the piRNA pathway (15, 24, 25) indicating a fundamental difference between the telomeric and conventional piRNA clusters. Therefore, the question of how the piRNA pathway affects telomere chromatin assembly in the germline is of particular interest.

Unique mapping of small RNA reads is the major source of information on the genomic origin of piRNAs, however, in the case of telomeres, it is technically challenging. Artificial sequences inserted into endogenous piRNA clusters serve as unique marks that allow for exploration of the highly repetitive genomic loci. Transgenic *Drosophila* strains carrying *P*-element copies in the terminal retrotransposon array have been identified and characterized (43). In contrast to the TAS that exert Polycomb group (PcG) protein-mediated silencing of transgenes inserted in these regions (44-46), the telomeric retrotransposon arrays show euchromatic characteristics and do not silence transgene reporters in somatic tissues, which in fact allowed selection of such transgenic strains (43). Therefore, based on their different ability to silence the integrated transgenes, two telomeric subdomains were defined within the *Drosophila* telomere in somatic tissues, namely transcriptionally active retrotransposon arrays and heterochromatic TAS (43, 44).

Taking an advantage of the telomere transgene model in combination with the experiments on endogenous telomeric elements, we investigated piRNA production and chromatin structure of the different telomeric loci in the ovaries of transgenic flies. It was shown that the production of telomere-specific piRNAs contributes significantly to chromatin structure and the expression of the studied telomeric regions in the germline. In contrast to somatic tissues, the TAS and *HeT-A—TART—TAHRE* arrays show similar chromatin structure and transcriptional status in the germline and can be related to the piRNA-producing domain. At the same time, we found that piRNA production is not similar between the transgenes integrated in different telomeric retrotransposons. Chromatin and cytological studies provide strong evidence that the telomeric piRNA clusters are highly sensitive to piRNA loss in contrast to the heterochromatic non-telomeric dual-strand piRNA clusters. Moreover, piRNA loss causes telomere translocation from the nuclear periphery towards the nuclear interior. These data, in combination with the previously observed discrepancy of telomeric and other dual-strand piRNA cluster response to the piRNA pathway disruption (15, 27, 47), suggest that a distinct type of piRNA cluster protects telomere integrity in the *Drosophila* germline.

## Results

### Transgenes located at different positions in telomeres produce small RNAs in *Drosophila* ovaries

It is well known that transgenes inserted within TAS produce abundant piRNAs and exert piRNA-mediated silencing of the complementary targets (48-51). However, the piRNA production ability of transgenes located within telomeric retrotransposon arrays has not been explored. In this study, we used four available transgenic EY strains on a *y^1^w^67c23^* (*yw*) strain background carrying the P{EPgy2} construct in the telomeric retrotransposon arrays (43). P{EPgy2} is a *P*-element-based vector containing *mini-white* and *yellow* genes. The transgene EY08176 was inserted into the GAG ORF of *HeT-A*-related *TAHRE* in the 2R chromosome. The transgenes EY00453 and EY00802 were integrated into the 3’ UTR of *TART-B1* of 3L while EY09966 was inserted into the *TART-C* of the 4^th^ chromosome. All *TART* insertions were in the promoter region located between the sense and antisense transcription start sites (39). All transgenes were mapped at between 12-23 kb distance from TAS (43). The orange eye colour of EY08176, EY00453 and EY00802 transgenic flies corresponds to the previously reported phenotype and indicates a high level of the *mini-white* reporter gene expression (43, 46). The EY03383 strain carries P{EPgy2} in the 2R TAS (43). The insertions in the TAS (EY03383) and in the telomere of the 4^th^ chromosome (EY09966) are silenced and demonstrate a white or variegated eye colour phenotype (Table 1). The euchromatic EY03241 transgene is used as a non-telomeric control. Insertion locations are shown schematically in Fig. 1a,b. DNA FISH on polytene chromosomes of salivary glands confirmed the telomeric localization of transgenes (Additional file 1: Figure S1).

**Figure 1.**
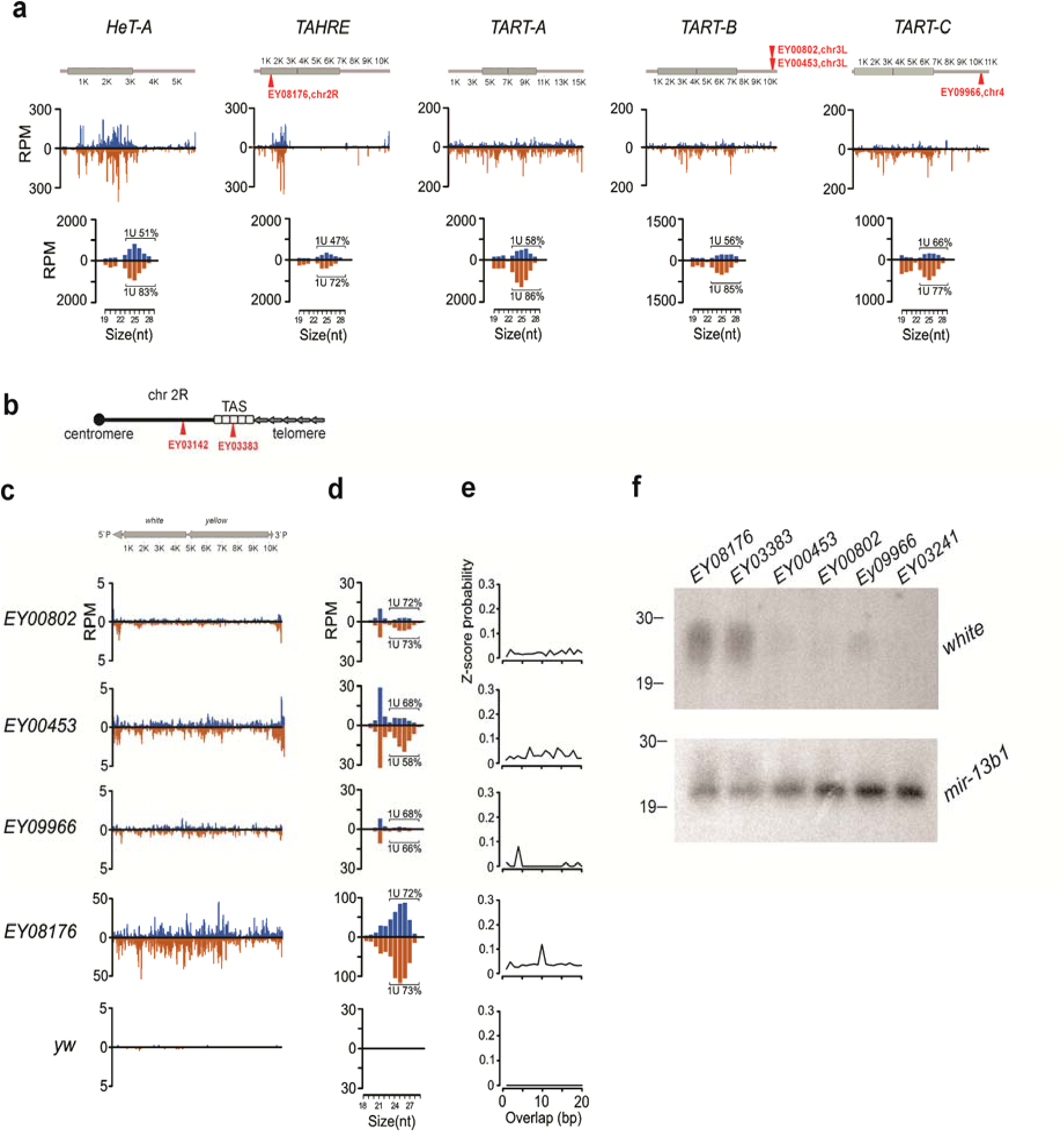
Generation of small RNAs by telomeric transgenes. (A) Schematic structure of telomeric elements is shown above. Insertion sites of transgenes are indicated as triangles situated above and below the schematics, which correspond to their genomic orientation. The profiles of small RNAs from ovaries of the *yw* strain are shown along the canonical sequences of *HeT-A, TAHRE, TART-B,* and *TART-C* telomeric retrotransposons. Normalized numbers of small RNAs (RPM, reads per million, 0-3 mismatches) in a 30-bp window were calculated. Length distribution of the telomeric element small RNAs is shown below. Percentages of reads having 1U are indicated for each strand (only 24-29-nt reads were considered). (B) Schematic of transgenic insertion sites in euchromatin and TAS of chromosome 2R. (C) Normalized numbers of small RNAs mapped to transgenic constructs (blue – sense; brown – antisense; no mismatches allowed). Schematic of the P{EPgy2} transgene is shown above. (D) Length distribution of transgenic small RNAs. Percentage of reads having 1U are indicated for each strand (only 24– 29-nt reads were considered). (E) Relative frequencies (Z-score) of 5’ overlap for sense and antisense 24–29-nt piRNAs (ping-pong signature). (F) Northern blot hybridization of the RNA isolated from the ovaries of EY08176, EY03383, EY00453, EY00802, EY09966, and EY03241 strains was performed with the *white* riboprobe to detect antisense piRNAs. Lower panel represents hybridization to *mir-13b1* microRNA. P^32^-labeled RNA oligonucleotides were used as size markers.

**Table 1.** Comparison of telomeric transgene properties in somatic tissues and in the germline

| strain [43] | insertion site | somatic tissues [43,44,46] |  | female germline |  |  |  |  |  |
| --- | --- | --- | --- | --- | --- | --- | --- | --- | --- |
|  |  | eye color | chromatin state | piRNA production | Rhi | HP1a | H3K9me3 | mini-white transcription | chromatin state |
| EY08176 | TAHRE Gag, chr.2R telomere | orange | active telomeric domain | +++ | +++ | +++ | +++ | + | dual-strand piRNA cluster |
| EY00802 | TART-B, 3' regulatory region, chr.3R telomere |  |  | + | + | +++ | +++ | + |  |
| EY00453 |  |  |  | + | + | +++ | +++ | + |  |
| EY09966 | TART-C, 3' regulatory region, chr.4 telomere | white | repressed telomeric domains | + | + | +++ | +++ | + |  |
| EY03383 | TAS, chr.2R | variegation |  | +++ | ++ | +++ | +++ | + |  |

+++ high level; + low level

We then asked whether the same transgenes inserted in the different positions of telomeric retrotransposon arrays produce a similar amount of piRNAs. To address this question, we sequenced the small RNAs from the ovaries of five telomeric transgenic strains and the EY03241 strain with a euchromatic insertion. Mapping of the small RNAs from the EY03241 strain to P{EPgy2} revealed a negligible amount of the transgenic small RNAs (Fig. 1c, Table S1).

Abundant endogenous *HeT-A, TAHRE* and *TART*-specific small RNAs are found in the *yw* and transgenic strains (Fig. 1a; Additional file 1: Figure S2), however, it is unclear, what the contribution of each particular telomeric element copy to the production of piRNAs is. Mapping of the small RNAs to telomeric transgenes revealed differences in the production of small RNAs (Fig. 1c), which may be attributed to piRNA production variations between the integration sites. The small RNAs are mapped to both genomic strands of the entire transgene EY08176 located within the *TAHRE* element. Most of the small RNAs mapping to the transgene are 24-29 nt long and demonstrate 5’ terminal uridine bias (1U bias), which is characteristic of piRNAs (Fig. 1d). We found the sense/antisense piRNA pairs (relative to transgene) overlap by 10 nt, which is a signature of the ping-pong piRNA amplification cycle (10, 18) (Fig. 1e). This small RNA profile strongly suggests that this transgene is integrated within the pre-existing piRNA cluster. The EY03383 transgene inserted in the dual-strand piRNA cluster within the 2R TAS produces abundant piRNAs from both genomic strands (Fig. 1c) similarly to the transgenes integrated into subtelomeric piRNA clusters on the X and 3R chromosomes (12, 47, 51).

The transgenes EY00453, EY00802 and EY09966 inserted in the *TART* elements within different chromosome arms produce much fewer small RNAs as compared to the EY08176 and EY03383, but more than the euchromatic EY03241 transgene (Fig. 1, Additional file 2: Table S1). A significant fraction of the small RNAs produced by the EY00453, EY00802, and EY09966 transgenes are 21-nt siRNAs; and no ping-pong signal was detected for the transgenic piRNAs demonstrating 1U-bias. Interestingly, the production of 21-nt RNAs is less variable between the telomeric transgenes than that of piRNAs (Additional file 2: Table S1). Unique mapping of the small RNAs to all telomeric transgenes revealed the piRNAs derived from the P-element fragments and linkers, confirming that the observed effects are transgene-specific (Additional file 1: Figure S3).

Northern blotting of the *white*-specific small RNAs from the ovaries of transgenic strains confirmed the presence of abundant small RNAs in the EY08176 and EY03383 strains (Fig. 1f, Additional file 1: Figure S4).

Thus, all telomeric transgenes can be considered as piRNA clusters, however, the piRNA-producing ability varies significantly between the transgenes integrated in different positions of the telomeric retrotransposon arrays (Table 1).

### HP1, Rhino and H3K9me3 associate with different telomeric transgenes

The piRNA-guided transcriptional silencing is mediated by the deposition of HP1 and H3K9me3 (14-17), whereas the germline-specific HP1 homolog Rhi serves as a chromatin marker of dual-strand piRNA clusters (23-26, 52). To answer the question as to whether these chromatin components are associated with different telomeric transgenes in *Drosophila* ovaries we performed chromatin immunoprecipitation (ChIP). The transcriptionally active *rp49* and *metRS-m* genes and the intergenic *60D* region were included in the analysis as negative controls. HP1 and H3K9me3 were considerably enriched in all studied transgenes (Fig. 2). As a positive control, we detected the Rhi enrichment at two regions of the *42AB* piRNA cluster (Fig. 2). Strong enrichment by Rhi was observed at two transgenic regions (5’ P-element arm and *mini-white*) in the EY08176 transgene, whereas a lower but statistically significant level of the Rhi binding was detected in the transgenes EY00453, EY00802, and EY09966 (Fig. 2; Additional file 1: Figure S5). The level of Rhi also varies at different positions of the *42AB*, which likely reflects the intrinsic heterogeneity of the chromatin structure in natural piRNA clusters.

**Figure 2.**
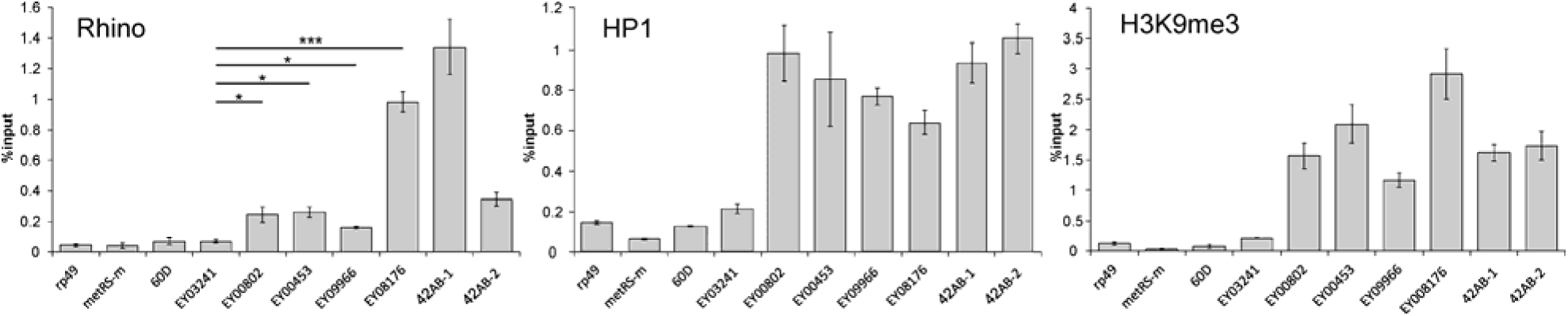
Chromatin components of the telomeric regions. HP1, H3K9me3, and Rhi occupancy at P{EPgy2} transgenes was estimated by ChIP-qPCR using primers corresponding to the 5’-P-element transgenic sequence. Two regions of the *42AB* piRNA cluster are enriched by all studied chromatin components. *rp49, metRS-m,* and *60D* regions are used as negative controls. Asterisks indicate statistically significant differences in Rhi enrichment relative to EY03241 (* P < 0.05 to 0.01, ** P < 0.01 to 0.001, *** P < 0.001, unpaired t-test). Difference in the HP1 binding between transgenes is statistically insignificant.

Judging by Rhi binding, which correlates with the ability to produce piRNAs, the telomeric transgenes belong to Rhi-dependent dual-strand piRNA clusters. Moreover, our data show that all telomeric transgenes, regardless of piRNA production rate and Rhi binding, associate with HP1 and H3K9me3 in *Drosophila* ovaries. These observations raise the question about the role of the piRNA pathway in deposition of HP1 and H3K9me3 crucial for the telomere functioning.

### piRNAs are required for the deposition and maintenance of HP1, Rhi, and H3K9me3 chromatin components at telomeric retrotransposon arrays in ovaries

HP1 and H3K9me3 are important components of telomeric chromatin involved in telomere length control in mammals (53). HP1 and H3K9me3 are also present in the *Drosophila* telomeres in somatic cells (44, 54, 55), however, the mechanisms underlying their deposition at the telomere are not clear and likely differ between the somatic and germline tissues. To study the role of the piRNA pathway in the deposition of HP1, H3K9me3, and Rhi at telomeres, we looked at the association of these proteins with telomeric transgenes and endogenous telomeric repeats following piRNA loss caused by depletion of the RNA helicase Spindle-E (SpnE) (3, 56). We demonstrated that the *spnE* mutation caused a considerable decrease in the association of HP1, Rhi, and H3K9me3 with the EY08176 telomeric transgene and with the endogenous *HeT-A* and *TART-A* elements accompanied by activation of their expression (Fig. 3; Additional file 1: Figure S6). This result is in agreement with the previously observed loss of H3K9me3 and HP1 from telomeric transposons upon piRNA pathway disruption (15, 42). In contrast to the telomeric regions, Rhi, HP1, and H3K9me3 are not displaced from the *42AB* locus and other dual-strand piRNA clusters in the ovaries of the *spnE* mutants (Fig.3). It is remarkable that chromatin of the EY00453 transgene inserted within the *TART* promoter is resistant to the piRNA loss caused by the *spnE* germline knockdown (GLKD) (Additional file 1: Figure S7) suggesting that the mechanism of chromatin maintenance at this particular site is different from other telomeric regions. Thus, ChIP data suggest that the piRNA pathway provides a germline–specific mechanism for the HP1, Rhi, and H3K9me3 deposition at different telomeric regions, and is essential for maintenance of this chromatin state during gametogenesis unlike the non-telomeric dual-strand piRNA clusters.

**Figure 3.**
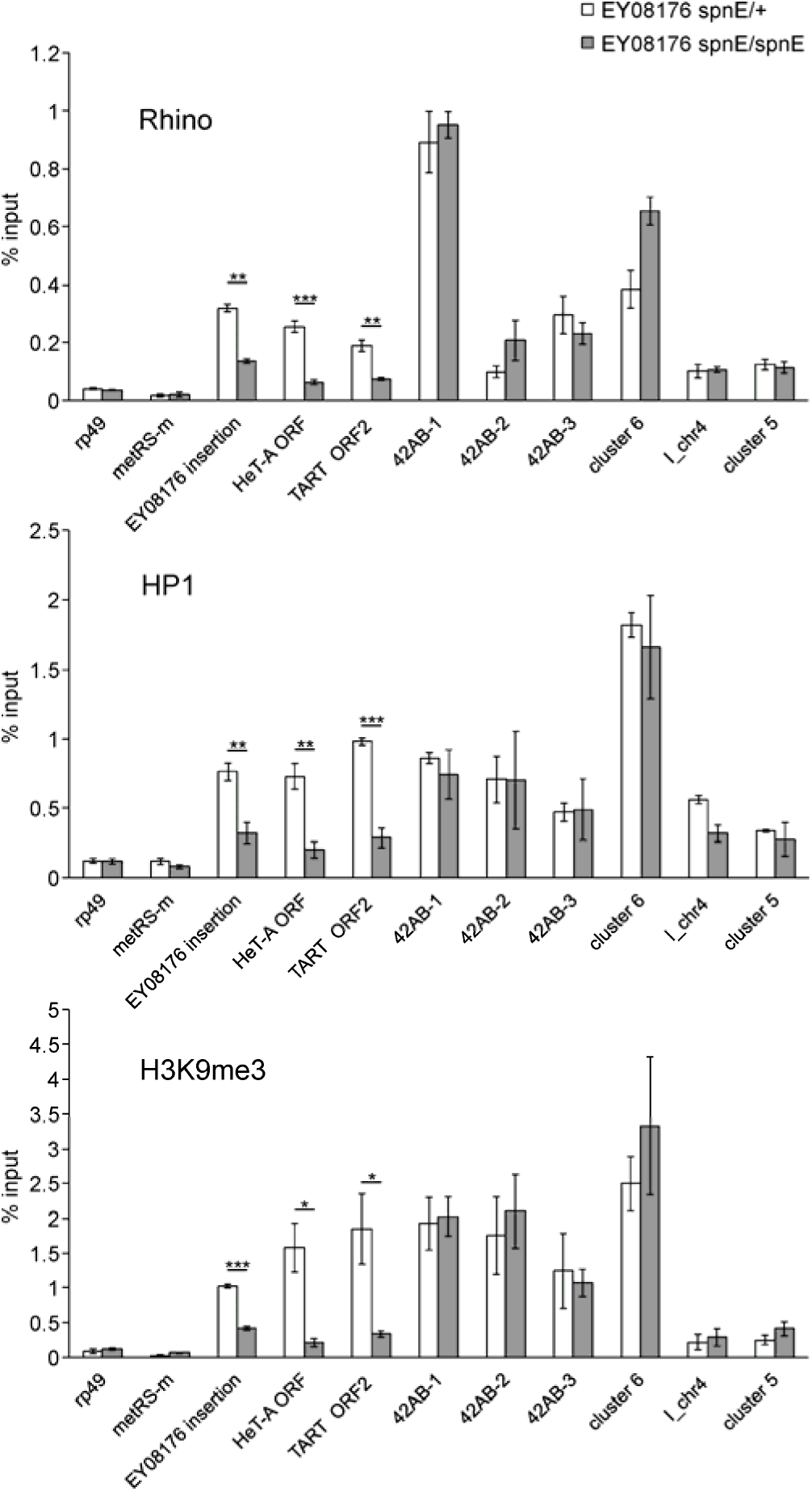
Role of the piRNA pathway in the deposition of HP1, Rhi, and H3K9me3 at telomeric transgenes in ovaries. ChIP-qPCR analysis of HP1, Rhi, and H3K9me3 enrichment at EY08176 telomeric transgene, endogenous *HeT-A, TART,* and several dual-strand piRNA clusters in ovaries of hetero- and trans-heterozygous (*spn-E^1^/spn-E^hls3987^) spindle-E* mutants. Asterisks indicate statistically significant differences in chromatin protein levels at indicated regions between *spnE*/+ and *spnE/spnE* (* P < 0.05 to 0.01, ** P < 0.01 to 0.001, *** P < 0.001, unpaired t-test).

### piRNAs are required for telomere localization at the nuclear periphery but are dispensable for telomere capping and clustering in the germline

To verify the Rhi association with endogenous telomeres in wild type ovaries and upon piRNA loss, we visualized *HeT-A* and *TART* using DNA FISH combined with Rhi immunostaining. In contrast to the giant polytene chromosomes of salivary glands, the chromatids of highly polyploid nurse cells are only partially conjugated allowing for the detection of numerous DNA FISH signals. In the ovaries of the *yw* strain, most of the *HeT-A* foci are clustered and overlapped with the largest Rhi foci forming rosette-like structures near to the nuclear envelope in the different *D. melanogaster* strains (Fig. 4a; Additional file 1: Figure S8a). We observed that the clustered *HeT-A* signals lose Rhi staining and are located toward the nuclear interior in the *spnE*, *piwi* and *zucchini (zuc)* piRNA pathway gene mutants (Fig. 4a; Additional file 1: Figure S8b). Positioning of the clustered *HeT-A* signals relative to the nuclear surface was estimated by 3D quantitative confocal image analysis of the *HeT-A* DNA FISH samples on the ovaries of control, *spnE,* and *piwi* mutant flies. It was found that the distance from the center of the *HeT-A* FISH signal to the nuclear periphery of nurse cells increased significantly in the *spnE* and *piwi* transheterozygous mutants as compared to heterozygous controls (Fig. 4b).

**Figure 4.**
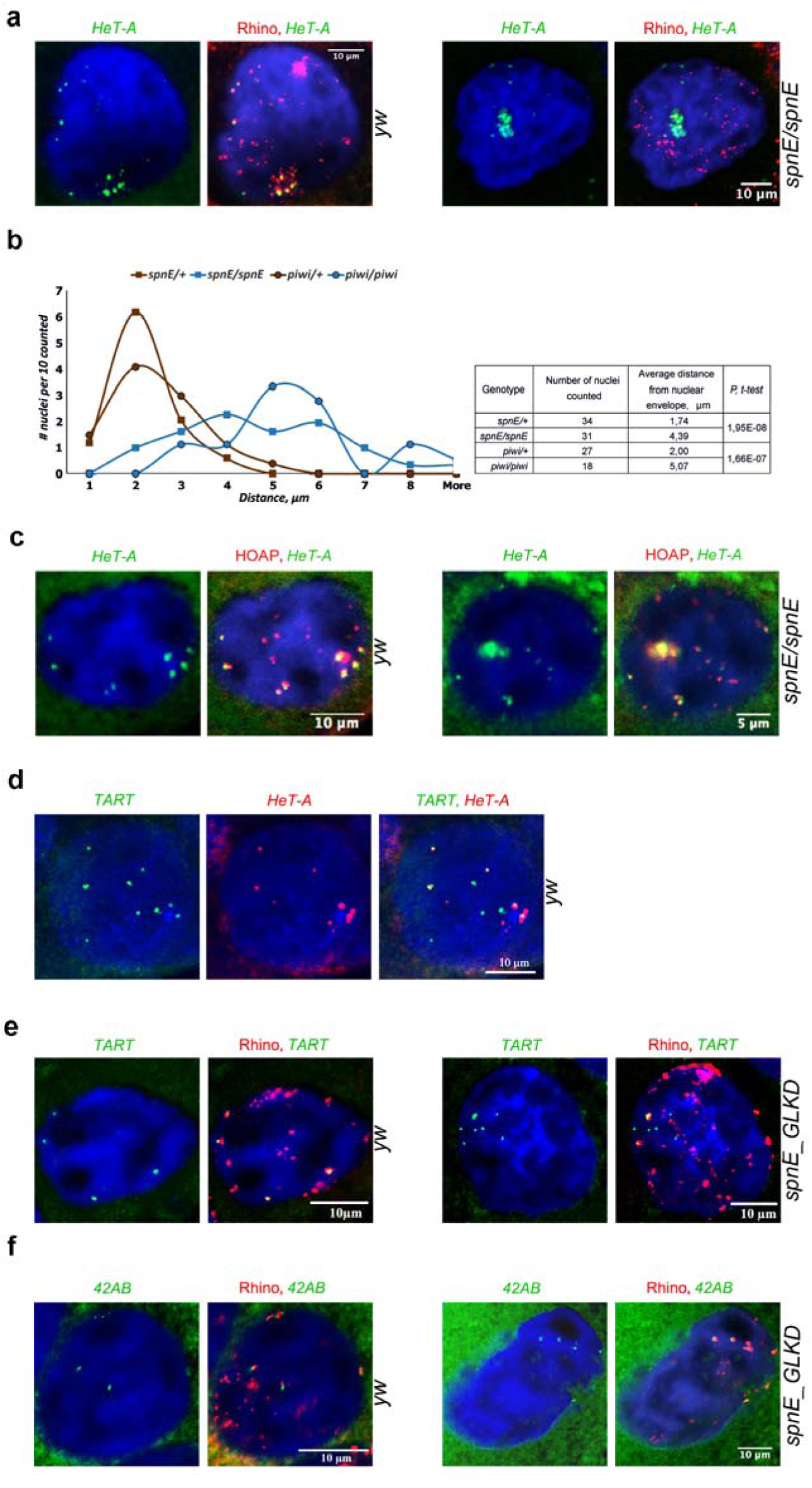
piRNAs are required for telomere localization at the nuclear periphery. **(A)** DNA FISH with *HeT-A* (green) combined with Rhi staining (red) was performed on ovaries of the *yw* strain and of the *spn-E^1^/spn-E^hls3987^* mutants. (B) Estimation of the positioning of clustered *HeT-A* signals relative to the nuclear surface of nurse cells by 3D quantitative confocal image analysis of *HeT-A* DNA FISH on ovaries of *spnE/+, spn-E^1^/spnE^hls3987^*, *piwi*/+, and *piwi^2^/piwi^Nt^* mutants. (C) piRNAs are dispensable for telomere capping and telomere clustering. DNA FISH with *HeT-A* probe combined with HOAP staining was performed on ovaries of the *yw* strain and of the *spn-E^1^/spn-E^hls3987^* mutants. (D) Double DNA FISH with *HeT-A* (red) and *TART* (green) probes was performed on ovaries of the *yw* strain. (E,F) piRNA pathway disruption causes loss of Rhi from *TART* but not from the *42AB* piRNA cluster. DNA FISH (green) with *TART* (E) or *42AB* (F) probes combined with Rhi staining (red) was performed on ovaries of the *yw* strain and of the *spn-E^1^/spn-E^hls3987^* mutants. DNA is stained with DAPI (blue). Nuclei of nurse cells from VIII-X stages of oogenesis are shown.

Next, we addressed the question about the role of piRNAs in the deposition of the protective capping complex at the chromosome ends in the germline. We performed the *HeT-A* DNA FISH combined with immunostaining of HOAP – the main component of the *Drosophila* telomere capping complex (57) – on ovaries of control flies and *spnE* mutants. HOAP extensively colocalizes with the clustered and individual *HeT-A* signals both in control and mutant nurse cell nuclei (Fig. 4c). Those HOAP signals that do not colocalize with the *HeT-A* most likely corresponded to telomeres lacking the full-length *HeT-A* copies since the *HeT-A* probe contains an ORF fragment. Previously, ChIP analysis has shown the reduction of *HeT-A* enrichment by HOAP in the *aubergine* and *armitage* but not in the *ago3* and *rhi* piRNA gene mutants (35). Thus, the HOAP loading at telomere ends appears to be mediated by specific piRNA pathway components (35) but not by piRNAs.

We suggested that the differences in chromatin structure and the ability to produce piRNAs among the transgenes integrated in different telomeric elements might be determined by the specific features of telomeric retroelements themselves. Using dual color DNA FISH with *HeT-A* and *TART* probes corresponding to their ORFs we showed that both *HeT-A* and *TART* had a different distribution in the nuclei of polyploid nurse cells. In contrast to the clustered *HeT-A* foci, a majority of the *TART* signals were independent and only a few of them colocalized with *HeT-A* (Fig.4d). Most likely, this pattern can be explained by the fact that the full-length *HeT-A* and *TART* are not present in all telomeres in the *yw* strain. In addition, *TART*-enriched telomeres seem to be not involved in telomere clustering in contrast to *HeT-A*-enriched telomeres. The *TART* DNA FISH combined with Rhi immunostaining demonstrates that the single *TART* signals colocalize with the small individual Rhi foci (Fig. 4e). This pattern is in agreement with the ChIP results showing that Rhi is deposited less in the *TART* and *TART* transgenes than in *HeT-A*. Colocalization of Rhi with *HeT-A* and *TART* DNA FISH signals decreases dramatically in the *spnE* mutants (Additional file 2: Table S2) in contrast to the *42AB* signals, which remain colocalized with Rhi (Fig. 4f).

Thus, piRNAs contribute significantly to the deposition of HP1, Rhi and H3K9me3 at the telomeric retrotransposon arrays and to the nuclear position of telomeres in the germline. However, they play only a minor role in the formation of telomere capping complex and telomere clustering.

### Comparison of subtelomeric chromatin in somatic and ovarian tissues

The *Drosophila* TAS regions consist of complex satellite-like repeats of 400–1800-bp in length and form heterochromatin domains that are able to induce silencing of transgenic constructs in somatic cells, a phenomenon known as telomeric position effect (58, 59). The TAS regions are enriched with H3K27me3 marks and bind Polycomb group proteins (44-46). In the germline, we observed HP1, Rhi, and H3K9me3 enrichment in the EY03383 subtelomeric transgene (Fig. 5a). ChIP using an anti-H3K27me3 antibody also revealed a high level of this chromatin mark at the transgene in ovaries of the EY03383 strain (Fig. 5b). This fact raises a question on how the different chromatin complexes coexist within the TAS regions.

**Figure 5.**
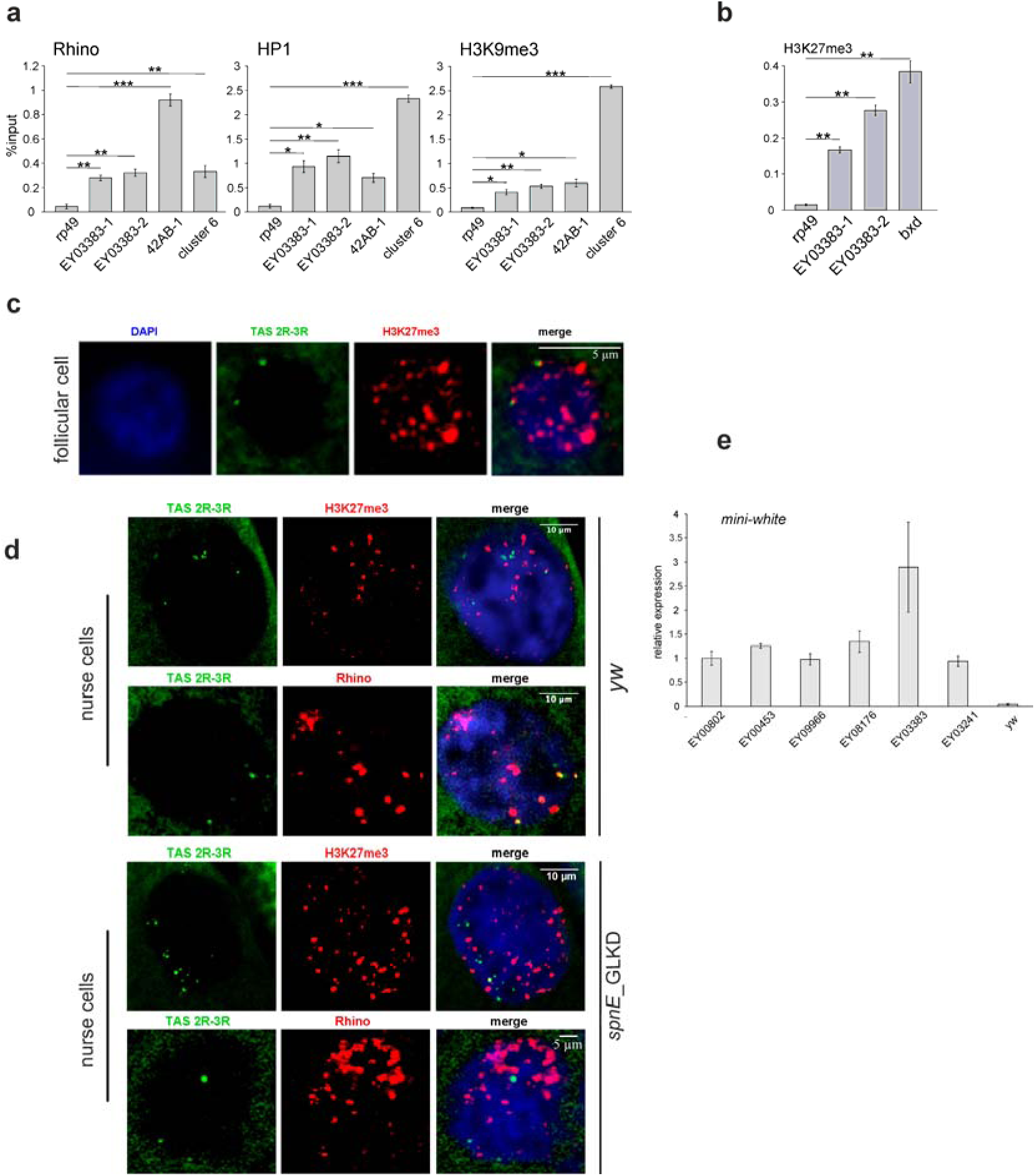
Comparison of subtelomeric chromatin structure in somatic and ovarian tissues. (A) HP1, H3K9me3, and Rhi occupancy at EY03383 transgene located in 2R TAS was estimated by ChIP-qPCR. *rp49* and *metRS-m* regions are used as negative controls. (B) ChIP-qPCR analysis of H3K27me3 enrichment at the subtelomeric transgene and *bxd* endogenous Polycomb-binding site on ovaries of the EY03383 strain. Asterisks indicate statistically significant differences in chromatin protein levels at the EY03383 relative to the *rp49* (* P < 0.05 to 0.01, ** P < 0.01 to 0.001, unpaired t-test). (C) DNA FISH with TAS 2R-3R probe (green) combined with H3K27me3 staining (red) was performed on ovaries of *yw* strain. The nucleus of a follicular cell is shown. (D) DNA FISH with TAS 2R-3R probe (green) combined with Rhi (red) or H3K27me3 (red) staining was performed on ovaries of the *yw* strain and of the *spn-E^1^/spn-E^hls3987^* mutants. Nuclei of nurse cells (stage VIII-X) are shown. (E) RT-qPCR analysis of the expression levels of transgenic *mini-white* in ovaries of transgenic strains. *white*-specific primers detect only transgenic transcripts because endogenous *white* is partially deleted.

An egg chamber comprises of an oocyte and fifteen nurse cells surrounded by somatic follicular cells. We suggested that TAS could recruit PcG proteins only in the somatic follicular cells. To visualize the relative position of TAS and proteins, we conducted DNA FISH combined with immunostaining on ovaries. The DNA probes corresponding to 2R-3R and 2L-3L TAS were used for FISH combined with a/Rhi and a/H3K27me3 immunostaining on ovaries of the *yw* strain. We observed a strong colocalization of the TAS probes with the H3K27me3 mark associated with Polycomb silencing in the nuclei of follicular cells (Fig. 5c, Additional file 1: Figure S9, Additional file 2: Table S3). On the contrary, the TAS signals show a much stronger colocalization with Rhi than with H3K27me3 staining in the nuclei of nurse cells (Fig. 5d, Additional file 2: Table S2, Table S3). We observed a loss of colocalization between the Rhi foci and TAS signals in the *spnE* mutants and upon *piwi* germline knockdown; overlap between the H3K27me3 staining and TAS probes was not considerably affected by *spnE* mutations in the nurse cell nuclei (Fig. 5d, Additional file 1: Figure S9a; Additional file 2: Table S2, Table S3). Thus, the PcG-dependent silencing of TAS is established in the ovarian somatic cells but not in the germline.

Next, we compared the expression level of telomeric transgenes in ovaries. We revealed that the steady-state RNA level of transgenic *mini-white* was similar in all the studied telomeric transgenic strains; and exceed the background signal detected in the *yw* strain in which the *white* locus was partially deleted (Fig. 5e). Simultaneously, active expression of the *mini-white* reporter was observed in the eyes of EY08176, EY00802, and EY00453 transgenic strains but not in the EY03383 and EY09966 strains. Thus, the transcriptional activity of transgenes located in different positions of the telomere is similar in the germline but differs considerably in the somatic tissues and appears to depend on the tissue-specific chromatin structure.

## Discussion

### piRNA production and Rhi binding differ along the telomeric region

To characterize telomeric piRNA clusters we integrated the data obtained from the analysis of endogenous telomeres as well as telomeric transgenes. The data on endogenous telomeric retrotransposons show that they produce piRNAs and associate with Rhi. However, the piRNA production by individual telomeric transgenes depends on the type of telomeric retrotransposon in which the transgene was inserted. The transgene located within the *TAHRE* produces considerably more piRNAs and shows stronger enrichment by Rhi than the transgenes located in the promoter region of *TART* elements. It is likely that the transgene integration *per se* in the *TART* regulatory region could interfere with *TART* promoter activity and reduce piRNA precursor read-through transcription. At the same time, Rhi immunostaining and *TART* FISH experiment also demonstrate that much less Rhi is deposited in *TART* than in *HeT-A* suggesting lower susceptibility of the *TART* elements to the engagement in piRNA production. Most likely, some features of the *TART* elements provide resistance to Rhi binding. Indeed, strong differences between the *HeT-A* and *TART* telomeric retrotransposons were observed in the genomic copy number, structure, patterns of transcription, and response to the piRNA pathway disruption (3, 29, 60, 61). *TART* transcripts are more stable (60), which can be explained by their role in providing reverse transcriptase (RT) for the transpositions of the main structural telomeric element *HeT-A* lacking RT. Therefore, one could suggest that the transcripts of full-length *TART* copies might be protected from piRNA processing to ensure encoding of the crucial enzyme for telomere elongation – *TART* RT.

Telomeric chromatin plays a pivotal role in telomere protection and maintenance. HP1 and H3K9me3 regulate capping, telomeric repeat silencing, and control of their transpositions onto chromosome ends (54, 55, 62). Interestingly, all the telomere insertions bind similar amounts of HP1 and H3K9me3 but strongly differ in Rhi association. Surprisingly, strong enrichment of the EY08176 transgene by Rhi, which recognizes the same H3K9me3 marks as HP1, does not abolish or significantly reduce HP1 binding compared to the insertions in *TART* elements indicating that Rhi and HP1 do not compete for binding sites at telomeric chromatin. Study of telomeric transgenes indicates that piRNA production and Rhi deposition are determined to a large extent by the type of telomeric retrotransposon into which they are inserted.

### Telomeric region represents distinct type of self-targeting dual-strand piRNA cluster

The piRNA sources and piRNA targets in the *Drosophila* germline are mainly represented by different genomic sequences; the piRNA clusters enriched with the damaged TE fragments provide the piRNA precursor transcripts, processed into piRNAs, that target active TEs (10, 15). The telomeric piRNA clusters have a dual nature and possess properties of both piRNA-clusters and piRNA-targets. It is well known that the piRNA targets are silenced at the transcriptional level through the assembly of repressive chromatin; loss of piRNAs causes a strong reduction in HP1 and H3K9me3 marks in complementary targets leading to their overexpression (14-16, 32, 42). However, piRNA loss fails to activate the germline piRNA cluster transcription and the switching from a repressive to an active chromatin state (15, 25, 27).

In-depth analysis of the telomeric piRNA clusters revealed strong differences in the chromatin dynamics between the telomeric and non-telomeric piRNA clusters. Using different approaches we demonstrated that piRNA pathway mutations induce the loss of HP1, H3K9me3, and Rhi from the telomeric transgene located in the *TAHRE—HeT-A* arrays as well as from endogenous telomeric retrotransposons in contrast to the other dual-strand piRNA clusters. It was shown that maternal and/or zygotic piRNAs were sufficient to induce formation of the repressive chromatin at non-telomeric piRNA clusters in early embryogenesis and that this state was maintained during germ cell development, even upon the loss of piRNAs at the later developmental stages (27). In contrast, piRNAs are required at all stages of germline development to maintain the telomere silencing. Accordingly, it was also reported that piRNA production of the *42AB* dual-strand piRNA cluster was far less sensitive to germline depletion of Rhi or HP1a than that of the subtelomeric piRNA clusters and transgenes located in this region (47). Thus, the chromatin dynamics of telomeric retrotransposons more resembles those of piRNA targets than those of piRNA clusters. At the same time, the telomeric regions bind Rhi and produce piRNA precursors, thus showing a relationship to the dual-strand piRNA clusters.

We believe that the fundamental difference between the Rhi-dependent telomeric and non-telomeric piRNA clusters is related to their different transcriptional regulation. Strong bidirectional promoters drive transcription of the telomeric retroelements (38-40). The loss of piRNAs causes activation of the promoters in telomeres resulting in a switching from a repressive to an active chromatin state (32, 40). In contrast, no discrete well-defined promoters were revealed within the heterochromatic non-telomeric piRNA clusters (37). Moreover, TATA box-binding protein (TBP)-related factor 2 (TRF2), previously described as a strong repressor of the *HeT-A/TAHRE* transcription, which is dispensable for *HeT-A* small RNA production (34), is required instead for transcription and piRNA production from the heterochromatic non-telomeric clusters (37). In fission yeast, high transcriptional activity at the siRNA target locus prevents heterochromatin assembly apparently through the displacement of the silencing complex (63). We found that the clustered *HeT-A* copies, normally positioned at the nuclear periphery were located more towards the nuclear center following the loss of piRNAs. We suggest that this process is induced by massive *HeT-A* overexpression and is related to the expression-dependent nuclear positioning phenomenon described by several groups (for review see (64)).

The telomere clustering near the nuclear periphery was observed in *Drosophila* somatic cells (65, 66). Telomeres are not clustered but do associate with the nuclear envelope in *Drosophila* oocytes at the pachytene stage of meiosis (67). We observe clustering of *HeT-A* DNA FISH signals near the nuclear periphery in the nuclei of polyploid nurse cells; however, it is unclear, which particular telomeres are involved in this clustering. The loss of piRNA affects the peripheral localization of telomeres in the germline; however, it does not affect telomere clustering or assembly of the telomere protection complex.

In addition to the telomeric regions, some recently transposed transcriptionally active TE copies inserted in euchromatin are related to the piRNA targets that also produce piRNAs (11). Strong reduction in H3K9me3 and Rhi association upon *piwi* depletion is observed for such TE copies (25). The main difference between the telomeric arrays and individual TE copies is that the latter are targeted by piRNAs mainly produced by other piRNA clusters or TE copies. We conclude that the telomeric piRNA clusters constitute a specific type of Rhi-dependent, actively transcribed piRNA clusters highly sensitive to the presence of piRNAs (Fig. 6).

**Figure 6.**
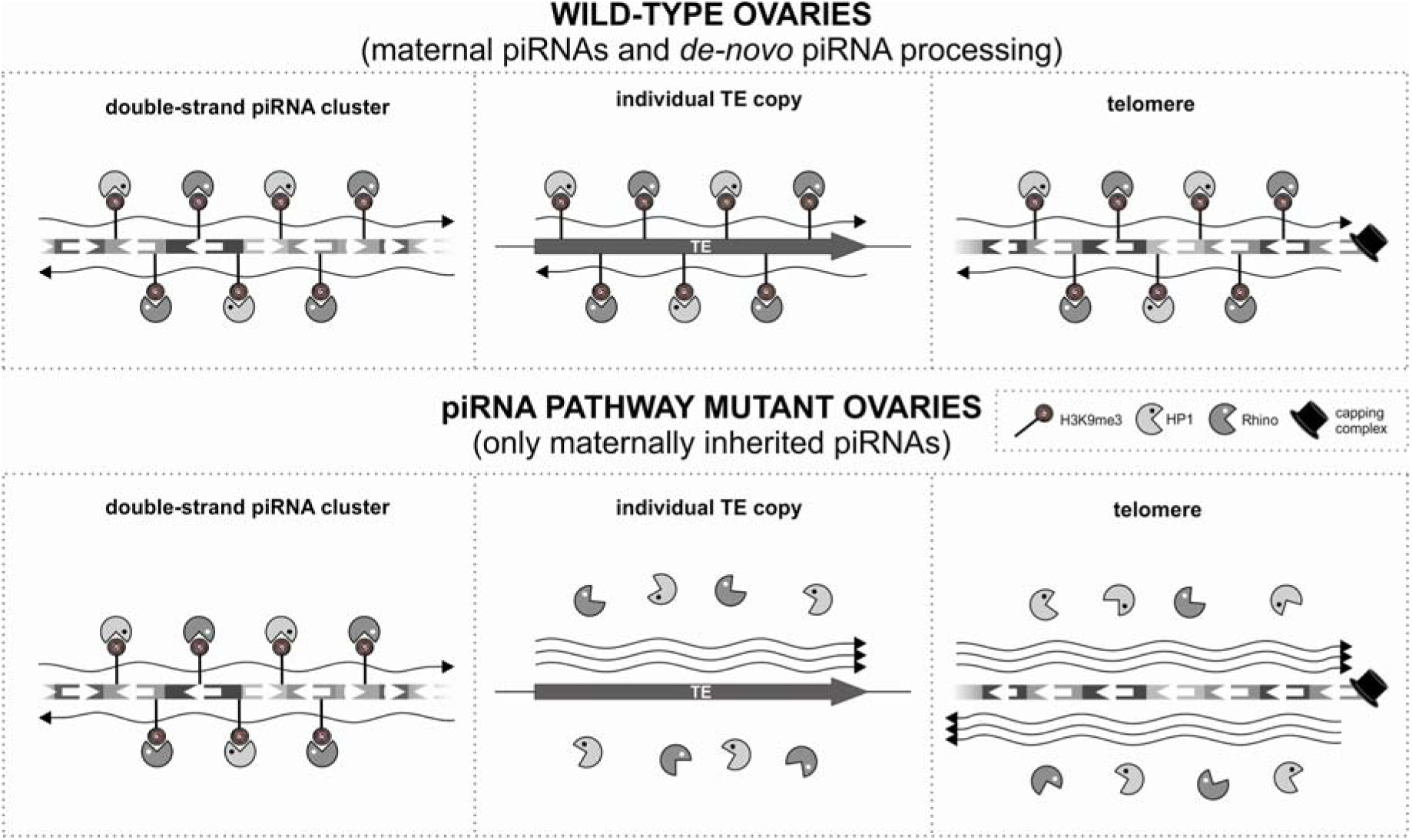
Telomeres represent a distinct type of the self-targeting dual-strand piRNA cluster. Schematic representation of three types of dual-strand piRNA clusters. Chromatin structure of canonical piRNA clusters is established by maternally inherited piRNAs but maintained by a piRNA-independent mechanism. On the contrary, piRNAs are strongly required for maintenance of chromatin state of telomeric and euchromatic TE-associated piRNA clusters during oogenesis. Only telomeric piRNA clusters produce piRNA precursors and piRNA targets at the same time. Assembly of telomere protection capping complex is not affected by piRNAs.

### Germline-specific chromatin structure of *Drosophila* telomeres

The comparison of expression and chromatin structure of the telomeric transgenes in ovaries and somatic tissues shows fundamental differences. Based on their ability to silence transgenes in somatic tissues, the TAS regions were defined as a heterochromatic domain, while the telomeric retrotransposon arrays were considered as a transcriptionally active subdomain (43, 44). Remarkably, the subtelomeric regions of diverse organisms consist of highly variable sequences that exert a silencing effect on transgenes integrated within these regions (68). Thus, the conserved silencing capacity of TAS is presumably important in the telomere functioning. We observe the similar chromatin properties of TAS and terminal *HeT-A—TART—TAHRE* arrays in the *Drosophila* germline. Both telomeric regions produce piRNAs, bind Rhi, and are expressed at a similar level (Table 1). Our data raise an intriguing question about the competition or developmentally regulated replacement of different chromatin complexes at TAS. The PcG protein binding sites were revealed in TAS repeats (45). Indeed, immunostaining and genetic analysis of the PcG protein mutants clearly demonstrates that the TAS zone serves as a platform for PcG protein-mediated chromatin assembly in somatic tissues (44, 45) and in ovarian somatic cells (Fig. 5). We suggest that initiation of piRNA precursor transcription in TAS displaces the PcG complexes or prevents their deposition in the germline. These tissue-specific silencing mechanisms have been observed by other groups; for example, the Polycomb repressive complexes were shown to silence transgenes carrying retrotransposon *Idefix* in somatic tissues but not in ovarian follicular cells (69). Interestingly, the retrotransposon *mdg1* copies marked by H3K27me3 in the ovarian somatic cells were not susceptible to piRNA-mediated transcriptional silencing (16). Our results in combination with the previous studies indicate that complex and competitive relationships between the various chromatin complexes define the chromatin structure of the genomic loci including telomeres, particularly in the developmental context.

## Methods

### Drosophila transgenic strains

Transgenic strains EY08176, EY00453, EY00802, EY09966, and EY03383 carrying the EPgy2 element and inserted within different telomeric regions were described previously (43) and were kindly provided by J. Mason. *Misy* natural strain was obtained from the collection of Institut de Genetique Humaine (CNRS), Montpellier, France. *P{EPgy2}Upf3^EY03241^* (stock #16558) was obtained from the Bloomington Drosophila Stock Centre. Strains bearing *spindle-E* (*spn-E*) mutations were *ru^1^ st^1^ spn-E^1^ e^1^ ca^1^*/*TM3*, *Sb^1^ e^s^* and *ru^1^ st^1^ spn-E^hls3987^ e^1^ ca^1^*/*TM3*, *Sb^1^ e^s^*. We used *piwi^2^* and *piwi^Nt^* alleles (70). *Zuc* mutants were *zuc^Hm27^/Df(2L)PRL* transheterozygous flies (71). GLKD (from “germline knockdown”) flies were F1 of the cross of two strains bearing construct with short hairpin (sh) RNA (spnE_sh, #103913, VDRC; piwi_sh, #101658, VDRC) and strain #25751 (*P{UAS-Dcr-2.D}1, w^1118^, P{GAL4-nos.NGT}40*, Bloomington Stock Center) providing GAL4 expression under the control of the germline-specific promoter of the *nanos (nos)* gene.

Fluorescence *in situ* hybridization (FISH) with polytene chromosomes was performed as previously described (72). A PCR fragment amplified using *white*-specific primers 5’-catgatcaagacatctaaaggc-3’ and 5’-gcaccgagcccgagttcaag-3’ was labeled with a DIG DNA labeling kit (Roche).

### RT-PCR analysis

RNA was isolated from the ovaries of 3-day-old females. cDNA was synthesized using random hexamers and SuperScriptII reverse transcriptase (Life Technologies). cDNA samples were analyzed by real-time quantitative PCR using SYTO-13 dye on a LightCycler96 (Roche). Values were averaged and normalized to the expression level of the ribosomal protein gene *rp49*. Standard error of mean (SEM) for two independent RNA samples was calculated. The primers used are listed in Additional file 2: Table S4.

### Small RNA library preparation and analysis

Small RNAs 19-29-nt in size from total ovarian RNA extracts were cloned as previously described (51). Libraries were barcoded according to Illumina TrueSeq Small RNA sample prep kit instructions and submitted for sequencing using the Illumina HiSeq-2000 sequencing system. After clipping the Illumina 3’-adapter sequence, small RNA reads that passed quality control and minimal length filter (>18nt) were mapped (allowing 0 mismatches) to the Drosophila melanogaster genome (Apr. 2006, BDGP assembly R5/dm3) or transgenes by bowtie (73). Small RNA libraries were normalized to 1 Mio sequenced reads. The plotting of size distributions, read coverage, and nucleotide biases were performed as described previously (13). Ovarian small RNA-seq data for *y^1^w^67c23^* and transgenic strains EY08176, EY00453, EY00802, EY09966, EY03383, and EY03241 were deposited at the Gene Expression Omnibus (GEO), accession number GSE98981.

### Chromatin immunoprecipitation

For every IP experiment ~200 pairs of ovaries were dissected. ChIP was performed according to the published procedure (74). Chromatin was immunoprecipitated with the following antibodies: anti-HP1a (Covance or C1A9 Developmental Studies Hybridoma Bank), anti-trimethyl-histone H3 Lys9 (Millipore), Rhi antiserum(40). Primers used in the study are listed in Additional file 2: Table S4. Quantitative PCR was conducted with a Light cycler 96 (Roche). Obtained values were normalized to input and compared with values at the *rp49* gene as a control genomic region. Standard error of mean (SEM) of triplicate PCR measurements for three-six biological replicates was calculated. Normalization of ChIP data on the *HeT-A* and *TART* copy number was performed using PCR on genomic DNA for each genotype. No substantial differences in the telomeric retrotransposon copy number were observed between *spn-E/+* and *spn-E/spn-E* flies.

### FISH and immunostaining

The combination of protein and DNA localization was performed according to the previously described procedure (72). Rabbit anti-H3K27me3 (Abcam) and rat anti-Rhi antibodies (40) were used. The probes used for DNA FISH were: *TART*, cloned fragment of *TART-A* ORF2 corresponding to 434-2683 nucleotides in GenBank sequence DMU02279; *HeT-A*, cloned fragment of *HeT-A* ORF corresponding to 1746 to 4421 nucleotides in GenBank sequence DMU 06920. *TART* probe was labeled using a DIG DNA labeling kit (Roche), *HeT-A* – by a Bio-Nick labeling system (Invitrogen). Probes corresponding to 2R-3R TAS, 2L-3L TAS, and *42AB* regions were PCR fragments obtained using primers listed in Additional File 2: Table S4 and labeled with a PCR DIG DNA labeling mix (Roche). To stain DNA, ovaries were incubated in PBS containing 0.5 µg/ml DAPI. Three biological replicas were obtained for each experiment. A Zeiss LSM 510 Meta confocal microscope was used for visualization. Confocal image z-stacks were generated with a slice step of 1.05 µM.

### Calculations of distance from the clustered *HeT-A* DNA FISH spots to the nuclear periphery

Calculations were performed using Imaris 7.4.2 software with manual segmentation of nuclei based on DAPI staining, automatic segmentation of *in situ* signal spots, and automatic calculation of a center of homogeneous mass corresponding to the main *HeT-A* cluster of FISH signals. FISH spot size is the diameter of a sphere encompassing all the spots in XY plane and Z position corresponding to the center of mass. The distance between the center of image masses and the nearest point on the nuclear surface was measured by increasing the radius of the sphere originating from the center of image masses until it intersected with the nuclear surface, and recording the radius as a distance. Independent two-sample t-test was used to compare hetero- and trans-heterozygous mutants.

### Northern blot of small RNAs

Northern blot analysis of small RNAs was performed as previously described (13). The *white* sense probe contained a cloned PCR fragment amplified using primers 5’-ctcacctatgcctggcacaatatg-3’ and 5’-attcagcagggtcgtctttccg-3’. Hybridization with P^32^ 5’-end-labelled oligonucleotide 5’-actcgtcaaaatggctgtgata-3’ complementary to the *miRNA-13b-1* was used as a loading control. The blots were visualized with a phosphorimager Typhoon FLA-9500 (Amersham). Northern blot quantification was performed using ImageJ.

## Additional files

**Additional file 1: Supplementary figures S1-S9.** (PDF 1494 kb)

**Figure S1.** Localization of telomeric transgenes

**Figure S2.** Profiles of telomeric retroelement small RNAs (related to Fig. 1a).

**Figure S3.** Generation of small RNAs by telomeric transgenes (related to Fig. 1c).

**Figure S4.** Quantification of Northern blots of small RNAs in transgenic strains (related to Fig. 1f).

**Figure S5.** Rhi and HP1 occupancy at telomeric transgenes (related to Fig. 2).

**Figure S6.** Expression of EY08176 telomeric transgene is increased in ovaries of the *spnE* mutants.

**Figure S7.** Chromatin structure of telomeric transgene EY00453 inserted in *TART* upon piRNA loss.

**Figure S8.** Nuclear localization of telomeres.

**Figure S9.** Subtelomeric chromatin in the germline (related to Fig. 5d).

**Additional file 2: Supplementary Tables.** (PDF 630 KB)

Table S1. Small RNA mapping to the telomeric and euchromatic transgenes.

Table S2 Colocalization of *HeT-A*, *TART* and TAS with Rhino.

Table S3 Colocalization of TAS with H3K27me3 in *Drosophila* ovaries. Table S4 Primers used in the study (5’-to-3’).

## Acknowledgements

Authors are grateful to J. Mason for transgenic *D. melanogaster* strains, Y. Schwartz for critical comments on the manuscript, Y. Rong for HOAP antibodies, and O. Olenkina for consultations on genetic crosses.

## Availability of data and materials

Ovarian small RNA-seq data for *y^1^w^67c23^* and transgenic strains EY08176, EY00453, EY00802, EY09966, ET03383 and EY03241 were deposited to the Gene Expression Omnibus (GEO), accession number GSE98981.

